# X-inactivation states of single cell transcriptomes reveal cellular phylogenies in human females

**DOI:** 10.1101/2022.11.10.515645

**Authors:** Alexander Predeus, Anna Arutyunyan, Laura Jardine, Chenqu Suo, Emma Dann, Regina Hoo, Martin Prete, Muzlifah Haniffa, Thomas J. Mitchell, Roser Vento-Tormo, Matthew D. Young

## Abstract

Human females undergo X-inactivation (Xi), whereby one copy of X is randomly inactivated early in development, then propagated through cell division. Because Xi state is inherited, its measurement in populations of cells encodes information about the phylogeny that created them and their relationships to other cells. We present a method, inactiveXX, to determine the Xi state of single cell transcriptomes, and demonstrate its accuracy using cancer and gold standard reference data. We apply inactiveXX to single cell transcriptomes from 190 human females, revealing that Xi in humans likely occurs around the 16 cell blastocyst stage and affects both embryonic and extra-embryonic tissues. We further find significant cell type specific variability in Xi skew, only detectable with cell type specific resolution, with certain cell types exhibiting strong population bottlenecks across tissues and disease state.

## Introduction

To ensure consistent dosage of X chromosome genes, most mammalian females inactivate one copy of their two X chromosomes. In humans, this X-inactivation (Xi) is established early in embryo development, with the inactivated copy of X randomly determined in each cell^1^. Once established, the inactivation status of the maternally or paternally derived X chromosome is stably inherited through subsequent rounds of cell division. Consequently, roughly half of somatic cells will have the maternally derived X inactivated. However, the precise ratio of maternal and paternal X-inactivation observed in any cells will depend on the relative contributions of the initially inactivated cells to that particular group of cells.

In extreme cases, where the contribution from a small number of the initially inactivated cells dominates, populations of somatic cells may exhibit strong X-inactivation skew. Pioneering studies used methylation status of the human androgen receptor gene *(AR)*, which has variable CAG repeats and is widely heterozygous, to detect the average Xi status of different tissues^2^. This work gave early insights into the clonal nature of cancer^3^, the size of progenitor populations^4^, and the identity of genes that escape Xi^5^. Despite these successes, measurement of Xi status using *AR* can only be used on a limited population, has an error profile that is difficult to remove^6^, and cannot easily be extended to measure specific cell types.

Single cell transcriptomics provides cell level readouts of gene expression on the X chromosome. The Xi status of a cell is encoded in this data via the specific alleles expressed at heterozygous single nucleotide polymorphisms (SNPs) on X. In this paper, we exploit this observation to develop a method, incativeXX, to determine the Xi status of individual cells from single cell transcriptomic data. We then demonstrate how these data can be used to understand phylogenies, population bottlenecks, and clonality of cells in a range of tissues from 190 human females, generating insights into the clonality of cells, cell type relationships, and the details of X-inactivation in humans.

## Evaluation of the method

To measure X-inactivation in single transcriptomes we developed a method, inactiveXX, which can infer the X-inactivation status of individual cells using single cell transcriptomic data alone. To achieve this, we consider 1,000 genomes^7^ SNPs with population frequency > 0.05 on X with evidence of both alleles across all cells, excluding regions known to escape X-inactivation^5^. We then count the number of reads supporting each allele in each single transcriptome. Using expectation maximisation we iteratively estimate the X-inactivation status of each cell (i.e., which copy of X is inactivated, the E-step) and the X chromosome genotype (i.e., phase the SNPs, the M-step) until convergence is reached. We repeat the EM fit 1,000 times using randomly chosen initial states, and use a deterministic annealing strategy to prevent local maxima^8^ **(Figure 1A)**.

**Figure 1.**
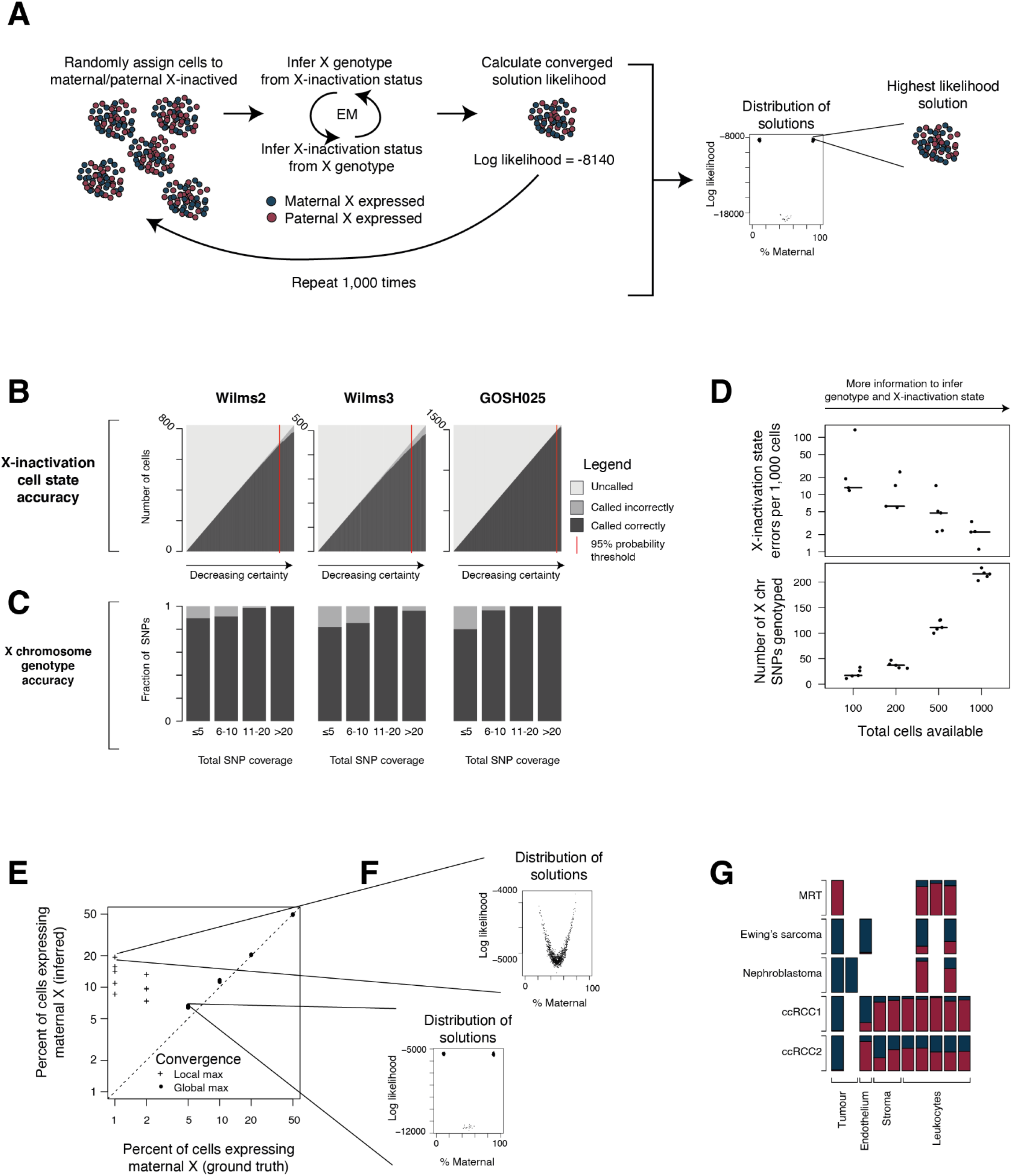
Method evaluation. **A - Overview of method:** Schematic overview of method to determine X-inactivation status. **B - Cell state accuracy on gold standard data:** Number of cells (y-axis) uncalled (light grey), called inaccurately (medium grey), and called accurately (dark grey) as a function of X-inactivation state probability (x-axis, decreasing to the right), for three individuals where parental and individual genomes define the ground truth. The 95% cut-off threshold is shown by a red line. **C - SNP accuracy on gold standard data:** Fraction of SNPs correctly phased (y-axis, colours as in **B**) as a function of total coverage across all cells (x-axis), for three individuals where parental and individual genomes define the ground truth. **D - Detection accuracy at different cell numbers:** Number of SNPs genotype (y-axis, bottom panel) and accuracy of X-inactivation status (y-axis, top panel) as a function of total cell number per individual (x-axis), where individuals are randomly generated 5 times for each cell number from gold standard data (**B,C**). Horizontal lines indicate median across random samples. **E - Detection accuracy at different X-inactivation skew:** X-inactivation skew detected across all cells (y-axis) compared to ground truth skew (x-axis), where individual data is generated by random sampling of gold standard data (**B,C**, dots). Dot shape indicates if the method believes it has (filled circles) or cannot (crosses) reach the global optimal solution. Dotted line indicates perfect correlation. **F - Example solution distributions for converged and non-converged fit:** Distribution of overall likelihood of solution (y-axis) as a function of total X-inactivation skew (x-axis) across all 1,000 random initialisations of method. The top plot shows an example where there is insufficient data to reach the global optimum solution, while the bottom plot shows an example where the global optimum has been reached. Note that solutions don’t cluster around any values in the non-converged plot, but instead lie on a continuum of ever increasing likelihood with more extreme X-inactivation skew values. **G - X-inactivation skew by cell types in cancer samples:** Fraction of cells in X-inactivation state (red/blue colour) for 5 different individual cancers (y-axis), by cell type (x-axis).

To evaluate the accuracy of this approach we first consider single cell transcriptomes of human females, for which individual and parental genomes are available^9,10^. Using the individual genomes we identify all heterozygous SNPs on X, phase them using parental genomes, and in each cell define the copy with the fewest supporting transcriptomic reads as inactivated. Applying our method and using a 95% confidence threshold, we determined the X-inactivation status of 90% of gold standard cells (range 86%-96%, **Figure 1B**), with 98% accuracy (range 95%-100%, **Figure 1B**), and correctly genotyped nearly all SNPs with >10 reads coverage (**Figure 1C**). We found our method could recover X-inactivation states of as few as 100 cells (90% accuracy, **Figure 1D**) and in populations with skews as extreme as 95-5 (**Figure 1E**). Importantly, we were able to identify when the true skew was more extreme than that detected by our method, but not recoverable with the data available (**Figure 1F**), preventing erroneous inference of X-inactivation status in extreme cases (e.g. skew of 100-0).

Next, we consider single cell transcriptomes from 5 cancers, where the cancer cell transcriptomes have been previously identified^9–12^. As cancers are derived from a single cell, cancer cells must have the same X-inactivation status. Across 5 samples, we found non-cancer cells have a range of X-inactivation skews, but cancer cells had on average 99% of cells in the same X-inactivation state (range 97.5-100%) (**Figure 1G**). Taken together, our method can accurately identify the X-inactivation status of individual cells from single cell transcriptomic data alone, even in cases of extreme X-inactivation skew or low cell number.

## Population dynamics and timing of X-inactivation

As Xi state is maintained through cell division, a group of cells X-inactivation pattern is determined by the Xi state of the founder cells (the cells present when Xi is determined) and the phylogeny that connects the observed cells to the founders (**Figure 2A**). Focusing first on a single individual, we inferred Xi states for 82,827 single cell transcriptomics from 6 fetal tissues^13^ (**Figure 2B**). Reasoning that cell type specific skew would be unbiased with respect to maternal/paternal X, we estimated the founder Xi skew as the median Xi skew across cell types (~80%). We then used a graph-based algorithm^14^ to identify groups of cells whose Xi skew significantly deviated from this value (**Figure 2C**). The local Xi skew matched the founder skew for almost all cell groups (96%, FDR 0.05), regardless of tissue or germ layer (endoderm, ectoderm, and mesoderm), consistent with all founder cells contributing roughly equally to all germ layers (**Figure 2D**). Only at the level of individual cell types did we see a significant shift in Xi skew for an appreciable fraction of cells (>10%), specifically in myelocytes, natural killer cells, erythroid cells, fibroblasts, and cycling epithelium (**Figure 2E**). Significant deviation from the founder Xi skew is evidence for a population bottleneck in these cells’ history. These cell types are enriched for those that are reactive or plausibly generated from locally seeded proliferation, which may be driving the observed Xi skew.

**Figure 2.**
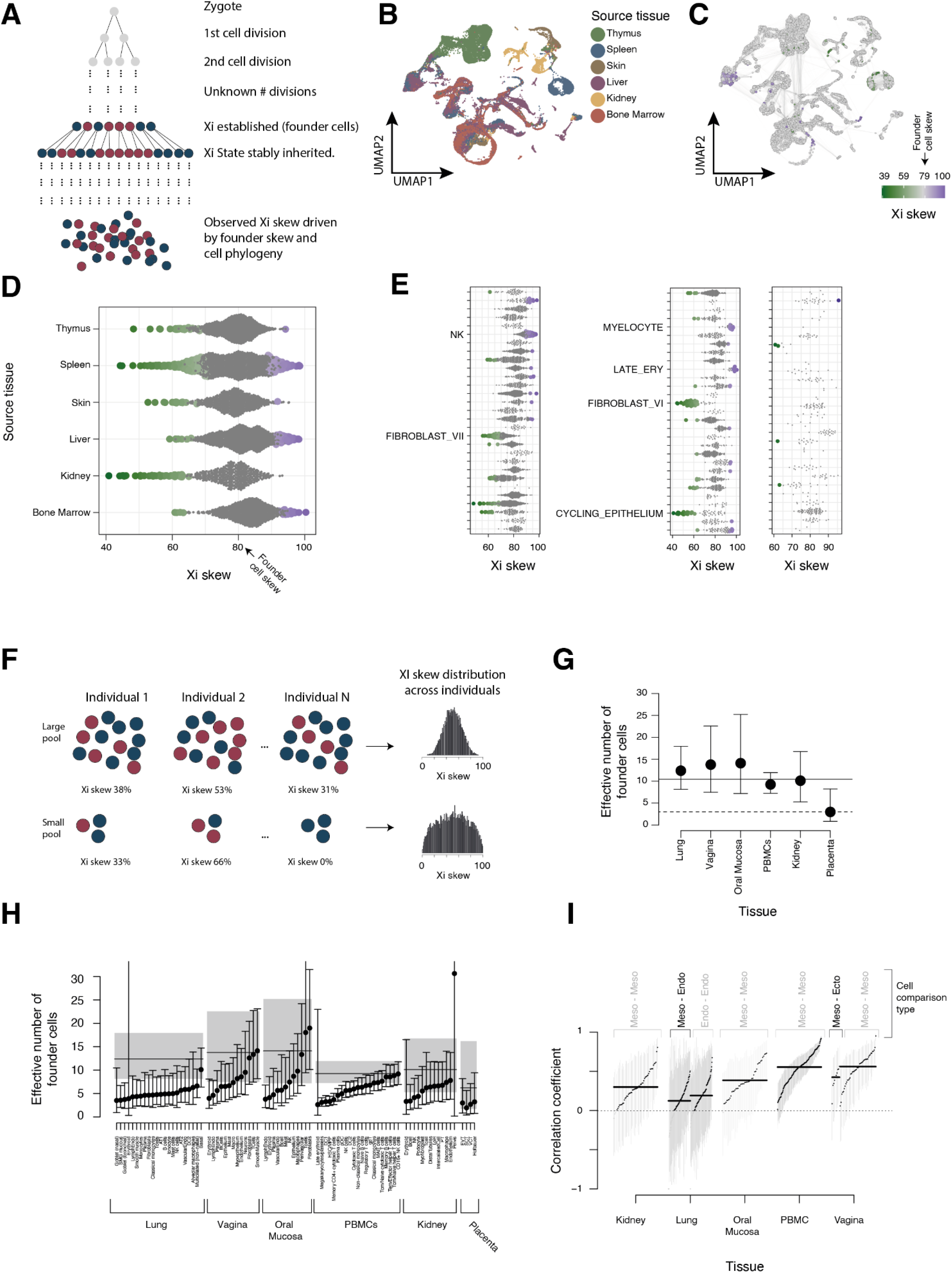
Population dynamics and timing of X-inactivation. **A - Illustration of connection between development, Xi, and observed cells:** Phylogeny showing key stages in X-inactivation. Cells preceding Xi are shown in grey. The 8 founder cells shown when Xi is determined are illustrative only, the true founder population size is not known. Once established the Xi status is inherited, with the Xi skew of the observed cells determined by the Xi status of the founder cells and how the observed cells are related to those founder cells (i.e., which founder cells contribute more or less). **B - UMAP showing tissue type of origin:** Reduced dimensional representation (UMAP) of fetal tissues from the same individual, with colour indicating tissue of origin. **C - UMAP showing significant regions:** UMAP showing neighbourhoods of cells from **B**, showing connectedness (line widths) and Xi state skew relative to the global average. Non-significant regions are shaded in grey, while significant ones are shaded by their Xi skew as given by the green-purple colour bar. **D - Significantly skewed neighbourhoods of cells by tissue:** Xi skew of neighbourhoods of cells from **B** (dots, x-axis), grouped by the dominant tissue type for that neighbourhood (y-axis). Neighbourhoods not significantly different from the global average are shown in grey, while significant regions are coloured as per the colour scheme in **C**, at three times the size. **E - Significantly skewed neighbourhoods of cells by cell type:** Xi skew of neighbourhoods of cells from **B** (dots, x-axis), grouped by the dominant cell type for that neighbourhood (y-axis), split across three panels to aid visibility. Neighbourhoods not significantly different from the global average are shown in grey, while significant regions are coloured as per the colour scheme in **C**, at three times the size. Cell type labels on the y-axis are shown for all cell types where >10% of neighbourhoods show significant Xi skew. **F - Connection between population size and X-inactivation skew distribution:** Illustration of how different primordial population sizes (rows) can give rise to different shaped Xi skew distributions (right) when measured across multiple individuals (columns) **G - Inferred founder population size by tissue:** Population size (y-axis) inferred from X-inactivation skew across multiple individuals in various tissues (x-axis). Points show point estimates, while error bars show the uncertainty due to cell and patient number. Horizontal lines show the overall median for embryonic (solid line) and extra-embryonic (dashed line) tissues. **H - Inferred founder population size at cell type resolution:** Population size (y-axis) inferred from X-inactivation skew across multiple individuals for various cell types (x-axis). Points show point estimates, while error bars capture uncertainty due to cell and patient number. Grey shaded regions show the tissue confidence intervals and horizontal line the tissue level mean, with tissues as labeled at the bottom of the x-axis. **I - Correlation of X-inactivation skew between cell types:** Pearson correlation coefficient (y-axis) between pairs of cell types (dots) within each tissue (x-axis labels), with 90% confidence intervals (vertical grey lines) quantifying uncertainty from cell and patient number. Horizontal lines show the population median, the dashed line shows no correlation, with values below 0 shown in grey. Comparisons are split by those within and between germ layers (top labels), with correlation coefficients for pairs of cell types from different (the same) germ layers shown in black (grey).

Xi skew may represent an individual specific event, or the result of a consistent biological process limiting the number of founder cells that contribute to a particular group of cells. To differentiate between the two, we inferred the Xi status from 6 tissue types, from 185 individuals, both with and without disease (**Table S1**). We reasoned that the spread of the cell or tissue Xi skew distribution across individuals is determined by the effective number of founder cells that give rise to that cell type of tissue^4^ (**Figure 2F**). This “effective number of contributing founder cells”, is a measure of the smallest population bottleneck that the observed cells are subject to, subsequent to the specification of random Xi. As such, it can never be larger than the number of cells present when Xi is determined, but can be smaller if founder cells are pre-specified to particular lineages, or a cell type/tissue undergoes a population bottleneck later in development.

Applying this analysis at the tissue level, to 5 tissues with at least 17 individuals (lung^15,16^, kidney^10,11,17^, oral mucosa^18^, vagina^19^, and peripheral blood^20^), we found a consistent value of ~10 founder cells (range 9-13) contributing to these tissues (**Figure 2G**). The consistency of this value across tissue types and disease states, and its consistency with previous estimates of founder population size^4^, suggest that when X-inactivation is determined in humans, approximately 10 cells exist that give rise to embryonic tissues. By contrast, we found the extra-embryonic placenta^21,22^ was derived from ~3 founder cells (95% confidence interval 2-7, **Figure 2G**). This finding is consistent with previous studies which found more extreme Xi skew in the placenta^23^. Furthermore, the ratio of embryonic to extra-embryonic founders (~3:10), matches the observation that roughly ⅔ of somatic cells are derived from just one of the cells following the first cell division^24–26^. Taken together, our data suggest that unlike in mice, where extra-embryonic tissue are not subject to random Xi, random Xi in humans is determined prior to the specification of extra-embryonic lineages, when the embryo is approximately 16 cells in size.

Next we estimated the effective number of contributing founder cells at the cell type level in each tissue (**Figure 2H**). In contrast to the tissue level analysis, we found considerable variation in effective founder number by cell type (**Figure 2H**). Unlike the embryonic/extra-embryonic tissue split, this difference cannot be driven by lineage segregation early in development, as a range of effective founder population sizes is evident across cell types known to be phylogenetically related (e.g. PBMCs **Figure 2H**). Instead, smaller effective founder number must be driven by later population bottlenecks, such as spatial lineage restriction (e.g. the calico cat effect at a cell type level) or local expansion of a limited pool of cells (e.g. tissue resident macrophages, vasculature derived from angiogenic expansion from a limited pool of tip cells). Of the cell types present across multiple tissues, we found consistently low effective founder numbers (i.e., small population bottlenecks) for erythroid, plasma, NK, and endothelial (vascular and lymphatic) cells, and bottlenecks in only certain contexts for fibroblasts, T cells, and macrophages. Overall, our data reveals that there is considerably more variability in Xi skew at the cell type level than tissue level, likely driven by cell specific phylogenies that are averaged out at the tissue level.

As the events that drive skewed X-inactivation are shared across cells with similar ancestry, we reasoned that correlation of Xi skew informs on cell type relatedness. Calculating the correlation of Xi skew between cell types revealed that the vast majority of cell types were positively correlated, a trend that held true across tissues and germ layers (**Figure 2I**). This is consistent with our pan-tissue analysis of a single individual (**Figure 2B-E**) and with all primordial cells contributing roughly equally to all three germ layers. However, we did find a lower correlation of cell types derived from different germ layers, compared to cell types from the same germ layer within the same tissue (**Figure 2I**). At the level of individual cell types, we found that the cell types with the strongest correlation within a tissue were those with a shared ancestry (e.g. tubular cells of the kidney **Supplementary Figure 1**).

## Discussion

We have presented inactiveXX, a new method to determine the X-inactivation status of individual cells from single cell transcriptomic data alone. Using cancer samples as positive controls and parental trios to define ground truth, we demonstrate that our method can accurately determine the X-inactivation status at single cell resolution in a wide range of conditions with very high accuracy.

Applying our method to a range of tissues and individuals suggested that X-inactivation in humans likely takes place prior to the specification of cells into embryonic and extra-embryonic, at the ~16 cell blastocyst stage. Furthermore, the inferred population size giving rise to embryonic and extra-embryonic tissues implies that ~¼-⅓ of cells of the blastocyst are restricted to extra-embryonic lineages, with the remaining cells giving rise to the embryo proper. This consistent picture at the tissue level hides significant variability at the level of individual cells, reflecting the ancestral relationships and clonal expansions of individual cell types. Correlating these cell level fluctuations across many individuals reveals the relatedness of individual cell lineages and that all cells of the early embryo contribute roughly equally to each germ layer. While the greatest power comes from cohorts of individuals, measuring the X-inactivation status of cells from the same individual can inform on the clonality and relatedness of individual groups of cells. We anticipate that the application of our method to large numbers of cells from the same individual and the same cell type across multiple individuals will deliver new insights into the clonality and developmental history of a wide range of cell types and tissues.

## Methods

### inactiveXX method

The inactiveXX method consists of four key steps:

1. Identify heterozygous SNPs on the X chromosome.
2. Generating allele specific counts for each cell at heterozygous SNPs.
3. Filter out low information SNPs (e.g. likely to escape Xi)
4. Fit EM model to estimate Xi state and genotype

Usually, the only data available are the single cell transcriptomes for which the Xi state is to be estimated. In this case steps 1 and 2 are performed simultaneously. However, when additional information is available, such as DNA sequencing from which heterozygous SNPs can be independently identified, each step may need to be performed separately.

#### Identification of heterozygous single nucleotide polymorphisms (SNPs)

Ideally, additional information, such as whole genome or exome sequencing would be used to identify heterozygous SNPs on the X chromosome. However, when this was not possible, we identified these from the single cell transcriptomic data.

To do this, we treated the single cell transcriptomic data as if it were derived from a single cell, and got the number of reads mapping to the reference and alternate alleles at sites of known variability on the X chromosome (1,000 genomes SNPs with population allele frequencies of 5% or more). Allele specific counts were generated using alleleCount (https://github.com/cancerit/alleleCount). At each site, we performed a binomial test against a null hypothesis of homozygosity of the other allele, with an error rate of 5%. We marked as heterozygous any site with p<0.05 for both alleles. That is, a site was considered heterozygous when there was a significant number of reads detected from both alleles, more than would be expected given an error rate of 5%.

#### Generation of allele specific counts

For each heterozygous SNP, we used alleleCount (https://github.com/cancerit/alleleCount) run in single cell mode using the -x flag, to generate counts at each site identified as heterozygous.

#### Filtering counts and SNPs

Before performing inference, we filtered counts on X to exclude those unlikely to be informative. By default, we excluded any SNP not lying in a known gene body (i.e., inter-genic), in a pseudo-autosomal region, or in evolutionary strata 1, 2, or 3, which have a high rate of Xi escape^5^.

We further filtered out any SNP that had evidence of expression of both alleles within a single cell, using a binomial test against a null hypothesis of only one allele being expressed and an error rate of 2%. We rejected any SNP with a p-value less than 0.1 as likely escaping Xi and so not informative for the inference step.

#### Inference of Xi state and X chromosome genotype

The input to the Xi inference step was cell specific counts for reference and alternate alleles, generated as described above. The inference step aims to simultaneously estimate which copy of X is inactive in each cell and which allele for each SNP (ref or alt) is on the maternally derived X chromosome using an expectation maximization approach. Note that “maternal” is used here just for convenience, it is only possible to infer which cells have the same X inactivated and which alleles belong to the same copy of X, not whether that copy of X is maternal or paternal (unless additional information such as parental DNA is available).

The inference followed the following steps:

1. Randomly set the Xi state for each cell and the X chromosome phasing for each SNP.
2. Set the deterministic annealing likelihood smoothing parameter, β, to 0.01.
3. In the E-step, estimate the Xi state probability from the current phasing as:

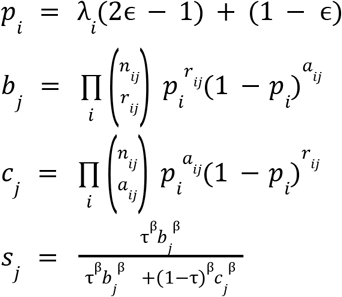

where *λ_i_* indicates if the maternal allele at SNP *i* is the reference 0, or alternate 1, ϵ is the sequencing error rate, *n_ij_* is the total counts at SNP *i* in cell *j*, *r_ij_* is the reference count at SNP *i* in cell *j, a_ij_*, is the alternate count at SNP *i* in cell *j*, τ is the fraction of cells in the maternal state (initialised to 0.5), and *s_j_*, is the probability that the cell *j* has the maternal X inactive. To determine when the EM algorithm has converged, we also calculate:

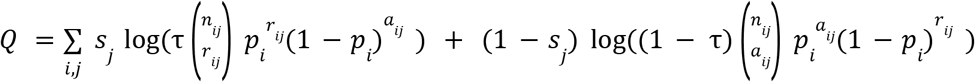
4. We then perform the M-step, where we estimate the SNP genotyping from the Xi states:

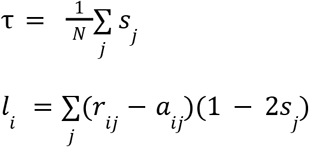

where *N* is the total number of cells and we then set *λ_i_* = 1 when *l_i_* > 0 and λ_i_ = 0 otherwise.
5. We iterate the E and M steps (steps 3. and 4.), until the difference in *Q* is less than tolerance (we use 1e-6) or 1000 iterations have passed.
6. We set β = min (1,1. 3 β) and repeat the EM fit (return to step 3.) until β = 1.
7. The final Xi states and X chromosome phasing (*λ_i_* and *s_j_*) are stored and the total likelihood (*Q*) is stored.
8. This fitting procedure is repeated 1,000 times (steps 1. −7.) and the results stored.
9. The best fit from the 1,000 random initialisations (fit with lowest *Q*) is taken as the accepted solution.

When there is sufficient data to accurately estimate the Xi state, the best fitting of the 1,000 fits will cluster around the same value in *Q* - τ space (up to reflection around τ = 0. 5). However, when the true value of τ is extreme, huge amounts of data (large cell numbers) are needed to accurately estimate it and there is no clear convergence amongst the 1,000 random starts.

We detected these globally non-converged solutions by requiring that the fractional difference between the best fitting value of τand the next 100 best solutions not exceed 5%. Samples where this was not the case were excluded from the analysis.

Those cells with state probabilities greater than 95% and non-significant number of counts that clash with the Xi state allocation (binomial test, 10% error rate and p-value cut-off 0.2) are declared to be “high-confidence”.

### Processing single cell data

#### Mapping data

For all single cell samples, raw nucleotide data was downloaded from the data provider and converted to fastq format using samtools^27^ v1.12 ‘collate’/’fastq’ commands where necessary. Following this, single cell RNA-sequencing experiments were aligned and quantified using the STARsolo pipeline^28^, according to the exact data type (3’/5’ 10x Chromium, inDrops, or Drop-seq). Wrapper scripts used for different technologies are available in the https://github.com/cellgeni/STARsolo repository. STAR aligner version 2.7.9a was used for all data processing.

Human reference genome and annotation exactly matching Cell Ranger 2020-A was prepared as described by 10x Genomics: https://support.10xgenomics.com/single-cell-gene-expression/software/release-notes/build#header. For 10x samples, the STARsolo command was optimized to generate the results maximally similar to Cell Ranger v6. Namely, “--soloUMIdedup 1MM_CR --soloCBmatchWLtype 1MM_multi_Nbase_pseudocounts --soloUMIfiltering MultiGeneUMI_CR --clipAdapterType CellRanger4 --outFilterScoreMin 30” were used to specify UMI collapsing, barcode collapsing, and read clipping algorithms. For paired-end 5’ 10x samples, options “--soloBarcodeMate 1 --clip5pNbases 39 0” were used to clip the adapter and perform paired-end alignment.

For inDrops samples, “--soloType CB_UMI_Complex --soloAdapterSequence GAGTGATTGCTTGTGACGCCTT --soloAdapterMismatchesNmax 3 --soloCBmatchWLtype 1MM --soloCBposition 0_0_2_-1 3_1_3_8 --soloUMIposition 3_9_3_14” options were specified; adapter sequence and whitelists varied between different versions of the protocol and were inferred directly from the fastq files.

For Drop-seq samples, “--soloType CB_UMI_Simple --soloCBwhitelist None --soloCBstart 1 --soloCBlen 12 --soloUMIstart 13 --soloUMIlen 8 --soloBarcodeReadLength 0” options were used for processing.

For cell filtering, the EmptyDrops algorithm employed in Cell Ranger v4 and above was invoked using “--soloCellFilter EmptyDrops_CR” options. Options “--soloFeatures Gene GeneFull Velocyto” were used to generate both exon-only and full length (pre-mRNA) gene counts, as well as RNA velocity output matrices. Coordinate-sorted BAM files were generated by STAR and indexed using samtools v1.12.

#### Demultiplexing mixed samples

A subset of samples contained multiple individuals mixed together in the same single cell sequencing run, necessitating computational separation of cells into individual specific groups. For fetal samples, we used the demultiplexing information provided^13^, and kept only those cells that were definitively fetal in origin (i.e., we excluded maternally derived cells). For other multiplexed samples, we used souporcell^29^, with k parameter equal to the number of expected multiplexed individuals to separate cells based on genotype, discarding those cells that could not be unambiguously genotyped.

#### Cell clustering and annotation

We took all barcodes identified by STARsolo^28^ as containing cell and produced log-normalised counts using the Seurat package^30^. Next we identified the top 2,000 most variable genes, which we used for principal component analysis. We retained the top 50 principal components and generated a two dimensional representation of the data using UMAP and clustered using a graph based clustering algorithm with a resolution parameter of 1.

Where mapping between barcodes and cell annotation were provided, we excluded all non-annotated cells and used the annotation provided. Where this was not the case, we annotated cells based on marker genes in the associated publication. For the two data-sets where neither published markers or pre-existing annotation were available (some kidney samples and PBMCs), we performed label prediction using previously trained models with CellTypist^31^, followed by manual inspection by experts in the relevant tissue.

### Validation of inactiveXX method

To validate the inactiveXX method, we used three samples with single cell transcriptomics, bulk DNA, and parental DNA to define a gold standard. To do this, we identified heterozygous SNPs in the bulk DNA, then phased them using the DNA from the parents, using the alleleIntegrator package^12^. This defined phased heterozygous SNPs on the X chromosome. We then used alleleCount to get the total number of reads mapping to the maternal and paternal copies of X in each cell (excluding standard low confidence regions such as the pseudo autosomal region) and defined the probability of each cell’s Xi state using the E-step equations (see above), with τ = 0. 5 and β = 1. For testing the accuracy of inactiveXX, we used only those cells whose Xi state could be allocated with high confidence (using the same definition as above) using the combined whole genome and transcriptomic data.

To measure the accuracy of inactiveXX as a function of individual level Xi skew and total number of cells, we repeatedly sub-sampled from the gold standard individual with the largest number of cells (GOSH025). To assess the effect of cell number, we sampled 100, 200, 500, and 1000 cells at random from this sample and re-calculated the accuracy with which transcriptomic data alone could recover the ground truth. For each number of cells, we repeated the random sampling 5 times.

To measure the accuracy with respect to the individual level Xi skew, we split cells into maternal and paternal Xi based on the ground truth data, then sub-sampled either the larger or smaller group (depending on the target individual level Xi fraction) to the number of cells necessary to create the desired individual level Xi skew. Once again, this random sampling was repeated 5 times.

### Xi skew in a single sample

To make inferences from Xi skew in a single sample, we first aimed to estimate the Xi skew in the founder cells. We assumed that while individual cell types may deviate from the founder Xi skew, there would be no systematic bias in the deviation. Provided this is true, then the founder Xi skew should agree closely with the mean (or median) of cell type specific Xi skews.

Having established an estimate of the founder Xi skew, we then aimed to test for statistically significant deviations from this value. We first generated overlapping, local neighborhoods of cells around representative cells in our data, from which the local Xi skew could be calculated, using the k nearest neighbour graph with miloR^14^. We sought to use the milo framework to test for shifts in Xi skew away from the provided founder skew. However, the milo framework uses a negative binomial model that requires multiple biological replicates to estimate the over-dispersion parameter for the model, which was not possible in the context of Xi skew. Instead, we used a binomial model (equivalent to setting the over-dispersion to zero) to calculate p-values, then used the graph based false discovery correction to correct for multiple hypothesis testing. We reasoned that this approach would err on the side of being permissive in identifying statistically significant deviations from the founder Xi skew. Finally, we marked any neighbourhood with FDR < 0.1 as significant.

### Effective founder population size analysis

To infer population size from Xi skew from the same cell or tissue type from multiple individuals, we assumed that the Xi skew variance across individuals would be given by a binomial distribution, with success probability 0.5, normalised by the number of cells. That is, a binomial distribution for n cells with *p* = 0.5, has variance σ^2^ = n/4. Normalising this by *n* produces a frequency distribution between 0 and 1, with σ^2^ = (π/4)/*n*^2^ = 4/*n*.

To account for the uncertainty in the estimate of Xi skew within each individual, we drew 1,000 samples from the posterior distribution of Xi skews for each individual, formed assuming a flat beta distribution prior. For each random sample across *n* individuals, we then calculated the effective population size as *n* = 4/σ^2^, where σ^2^ represents a random sample from the sampling distribution of the variance across *n* individuals. The quantiles of the resulting 1,000 values for *n* then define the uncertainty in our estimate of the population size, accounting for both the uncertainty due to the number of cells used to estimate the Xi skew in each individual, and the uncertainty due to the number of individuals used to estimate the variance in Xi skew across individuals.

When applying this analysis to the placenta, we considered only those cells that were definitively of extra-embryonic origins (trophoblasts).

### Correlation of cell types

To estimate the correlation of Xi skew between cell types, we calculated the Xi skew of each cell type and its uncertainty as described above. Given two vectors of Xi skews, one per cell type, of length n, representing *n* individuals, we then calculated a pearson correlation coefficient. To quantify the uncertainty in this estimate, we again drew random samples from the sampling distribution of the correlation and took quantiles of 1,000 random samples to define the uncertainty. We took samples from the exact distribution in this paper, but also provide a computationally faster implementation based on the Fisher transformation as part of the inactiveXX package.

## Data availability

Data were downloaded from the HCA data portal (https://data.humancellatlas.org/), using the links specified in **Table S1**, or for non-HCA samples, from the reference provided in the associated publication listed in **Table S1**. Additionally, we provide the Xi state probabilities, annotation, and Xi state calls for all cells processed in this sample in **Table S2**.

## Code availability

The inactiveXX method, along with documentation, is available as an R package (https://github.com/constantAmateur/inactiveXX). The code used to process the samples in this paper and produce the resulting plots is also made available (https://github.com/constantAmateur/XiPaperCode).

## Author contributions

M.D.Y. conceived the study, wrote the method, performed the analysis, and wrote the manuscript. T.J.M assisted with the analysis. A.P. downloaded, processed, and mapped the single cell data. A.A., L.J., C.S., E.D., M.P., and M.D.Y, performed and evaluated the tissue specific annotations. R.H. performed experiments for placental data. M.H. contributed to understanding the fetal data. R.V-T. provided guidance regarding extra-embryonic tissues and contributed to understanding the placental data.

## Supplementary Figures

**Figure S1.**
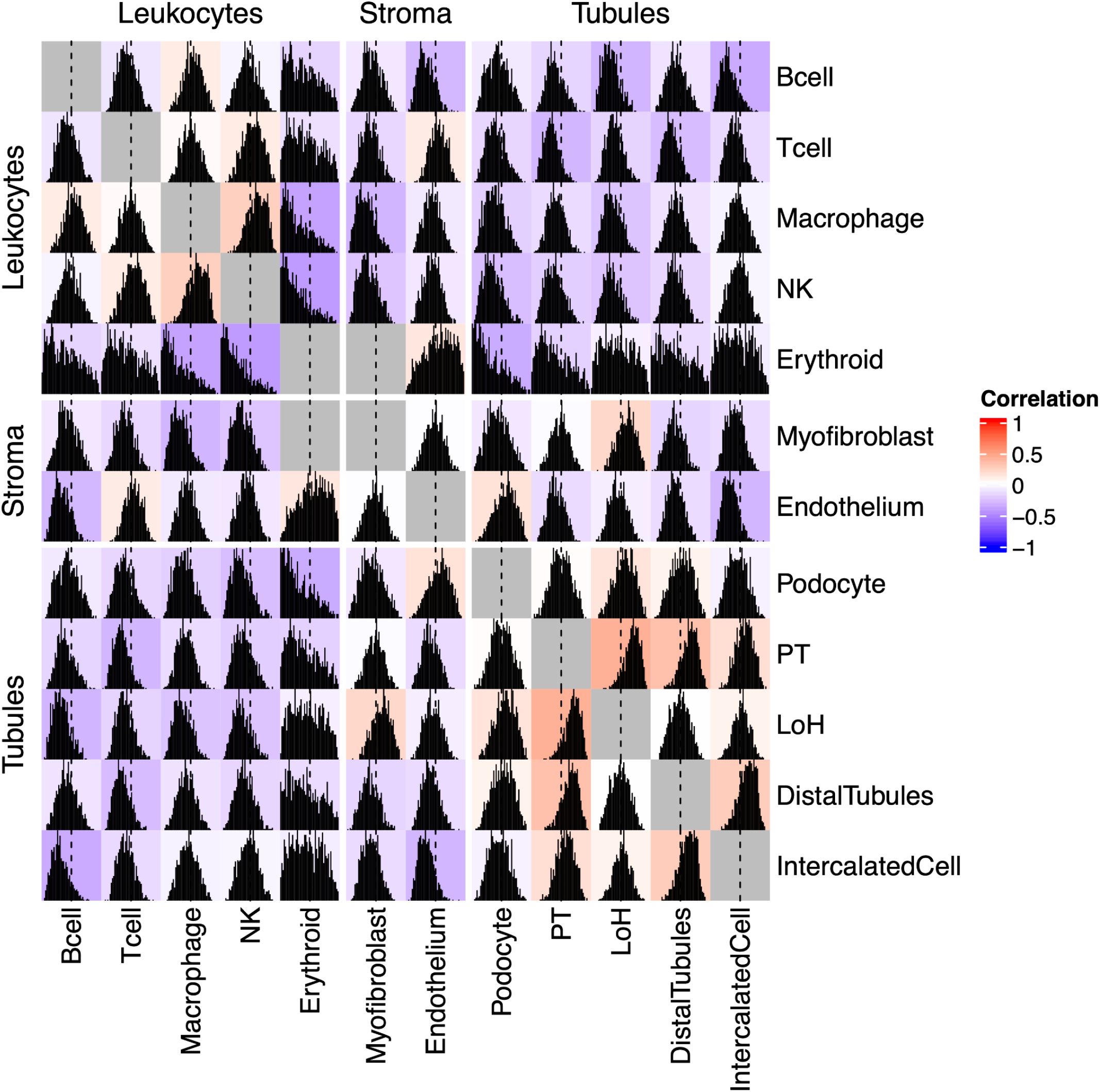
Correlation of Xi skew amongst cell types of the kidney. Correlation of Xi skew across multiple individuals, relative to the within individual cell type average (i.e., the founder Xi skew), for cell types of the kidney (bottom and right labels), grouped into cell type categories (left and top labels). For each comparison, the average correlation coefficient (colour value), posterior probability distribution for the correlation coefficient (histogram), and 0 correlation value (vertical dashed line) are shown for each square.

## Supplementary Tables

**Table S1 - Meta-data relating to individuals present in this study**

For each individual with data present in this study, this table provides information about the data type, tissue, individual age, disease status, and data source.

**Table S2 - Meta-data relating to individual cells present in this study**

For each cell used in this study, this table provides information about where it was derived from, which cell type it was annotated as, its inferred Xi state, and the gold standard Xi state (where available).

